# Isolation and Lipidomic Profiling of Neuronal Lipid Droplets to Unveil the Lipid Landscape of Neurodegenerative Disorders

**DOI:** 10.1101/2023.12.13.571527

**Authors:** Mukesh Kumar, Rachel McAllister, Souvik Roy, Justin A Knapp, Kallol Gupta, Timothy A. Ryan

**Author notes:** These authors contributed equally (Mukesh Kumar, Rachel McAllister).

## Abstract

Recent advances have expanded the role of lipid droplets (LDs) beyond passive lipid storage, implicating their involvement in various metabolic processes across mammalian tissues. Neuronal LDs, whose existence has long been debated, have now been identified in several neural structures, raising questions about their contribution to neurodegenerative disorders. Elucidating the specific chemical makeup of these organelles within neurons is critical for understanding their role in neural pathologies. However, the inherent submicron size and low abundance in neurons have posed challenges in their isolation; and physical and biochemical characterization. Here, we present our protocol which overcomes this obstacle: we blocked neuron-specific triglyceride lipase to enlarge LDs, rendering them amenable to floatation in a sucrose gradient. We further employed an improved methodology of lipidomic analysis on the LDs purified from cultured primary neurons, offering insights into their unique lipid composition. This protocol can be implemented by researchers with basic expertise in neuronal culture, imaging, ultracentrifuge handling, and LC-MS analysis. The complete workflow requires 3-4 weeks, from neuronal culture to lipidomics analysis. Future integration of this method with high-throughput techniques holds promises for uncovering disease-specific alterations in lipid metabolism, providing new avenues for potential therapeutic interventions.

**Key points:**

- This protocol describes a robust workflow for pharmacological induction, purification, and characterization of mature lipid droplets from primary cortical neurons without any detectable cytosolic membrane carryover.
- LC-MS analysis of neuronal lipid droplets can reveal lipids previously uncharacterized by other methods, providing a platform to integrate high-throughput proteomics and other techniques for probing neurodegenerative disease linked metabolic alterations in neurons.

**Key reference:** Kumar, M. *et al.* Triglycerides are an important fuel reserve for synapse function in the brain. *Nat Metab* **7**, 1392-1403 (2025). https://doi.org:10.1038/s42255-025-01321-x

**Editorial summary:** This protocol describes a robust workflow for pharmacological induction, purification, and characterization of mature lipid droplets from primary cortical neurons without any detectable cytosolic membrane carryover.

**Proposed teaser:** Isolation and lipidomic profiling of neuronal lipid droplets

## INTRODUCTION

Lipid droplets (LDs) were traditionally viewed as passive reservoirs of esterified lipids in adipocytes. Recent advancements in imaging and biochemical techniques have, however, detected the presence of LDs in most mammalian tissues, and they are now recognized as dynamic organelles involved in multiple aspects of cellular functions, including energy production, membrane synthesis, and signaling pathways (see Farese *et al.*^1^ and Olzmann *et al.*^2^ for comprehensive reviews)^2,3^. Recent studies have also established lipid dysregulation and excessive LD accumulation as hallmarks of metabolic disorders such as obesity^4^, diabetes^5^, atherosclerosis^6^ and fatty liver disease^7^. This suggests that the LDs are more than reservoirs of triglycerides (TG). In non-neuronal tissues, they are enriched in cholesteryl esters, ether lipids, signaling phospholipids, bioactive intermediates and bioactive proteins that can modulate organelle crosstalk, and stress response^8,9^.

Neurons, with their complex morphology and high energy demand, are not exempt from the importance of lipid metabolism, and lipids are increasingly gaining attention as a potential key player in the functioning and malfunctioning of neurons. Contrary to the prevalent belief that LDs are non-existent in neurons, they have been detected in motor neurons of drosophila^10^, axons of *Aplysia*^11^, cerebral cortex of mice brain^12,13^, and neuroblastoma-derived cell lines^14^. Despite multiple reports of lipid accumulation in neurodegenerative disorders (ND)^12,15,16^, the proteo-lipid identity of this organelle in neurons has remained undefined, in part because of the lack of a reproducible neuron-specific LD isolation protocol.

Neurodegenerative diseases, like Alzheimer’s disease, Parkinson’s disease, and Hereditary spastic paraplegia are multifactorial disorders that share a common theme of progressive degeneration of neurons, but their precise etiology remains elusive. Interestingly, abnormal accumulation of LDs in these disorders^12,15–17^ has raised the question of whether the impaired TG mobilization—across cell types in the brain and within intracellular organelles—contributes to neuronal dysfunction and deterioration in vulnerable brain regions. Evidence suggests that neuronal lipid droplets (NLDs) might serve as the site for accumulation of both neuroprotective and toxic lipid, and protein species^18,19^. Moreover, LDs play a crucial role in cellular communication and protection against inflammation^20^.

Recent studies further point to a tight coupling between lipid metabolism and bioenergetics in the brain. Astrocytic oxidative phosphorylation (OxPhos) is essential for hydrolysis of fatty acids (FAs) and maintenance of lipid homeostasis in the brain. Abnormal OxPhos in astrocytes causes LD accumulation followed by demyelination and neurodegeneration^21^. A related line of evidence comes from the patients with hereditary spastic paraplegia type 54 (HSP54, also known as SPG54), a rare autosomal-recessive neurodegenerative disorder caused by mutations in DDHD2 lipase gene. Cerebral proton magnetic resonance spectroscopy (MRS) of the patients’ brains manifest abnormal spectra with a lipid peak consistent with altered brain lipid metabolism^22,23^. Moreover, genetic silencing or pharmacological inhibition of DDHD2 lipase in mice causes massive accumulation of TGs specifically in the brain^12^. Although the precise molecular mechanisms linking TG metabolism to neurodegeneration remain unknown, current evidence supports a close relationship between lipids and neuronal energy metabolism. We recently discovered that DDHD2 controls the flux of FAs to mitochondria to support synaptic function under hypometabolic conditions^24,25^. Intriguingly, electrical activity triggers the transfer of FAs from triglyceride-rich LDs to mitochondria for ATP synthesis in dissociated hippocampal neurons^24^. Sortilin- ApoE3 signaling enables neurons to use long-chain FAs as an alternate fuel^26^. In parallel, work in *Drosophila* shows that neuronal FA oxidation is required to fuel memory formation after intensive learning^27^.

### BOX1 Ultrastructure of lipid droplets

Lipid droplets arise by phase-separation of esterified lipids on the outer leaflet of the endoplasmic reticulum. When cells accumulate an excess of fatty acids, neutral lipids coalesce at the ER membrane to form nascent LDs28. The nascent LDs then bud off towards the cytosolic side, forming a core of esterified lipids (e.g., triglyceride ester, cholesterol esters) encased by a monolayer of phospholipids and enzymes involved in lipid metabolism. EM microscopy reveals that the LDs are often associated through lipidic-bridges to the ER or/and mitochondria depending on the metabolic state of the cells29. These lipid-bridges function as active channels for the exchange of class I LD proteins and as well as lipids between organelles. A subset of proteins (class II) is also recruited to LDs directly from the cytosol through a lipid anchor motif30. In neurons, LDs are localized both in the soma and extended processes, often in close apposition to mitochondria24—underscoring their role in inter-organellar communication and functional integration for synaptic energy homeostasis.

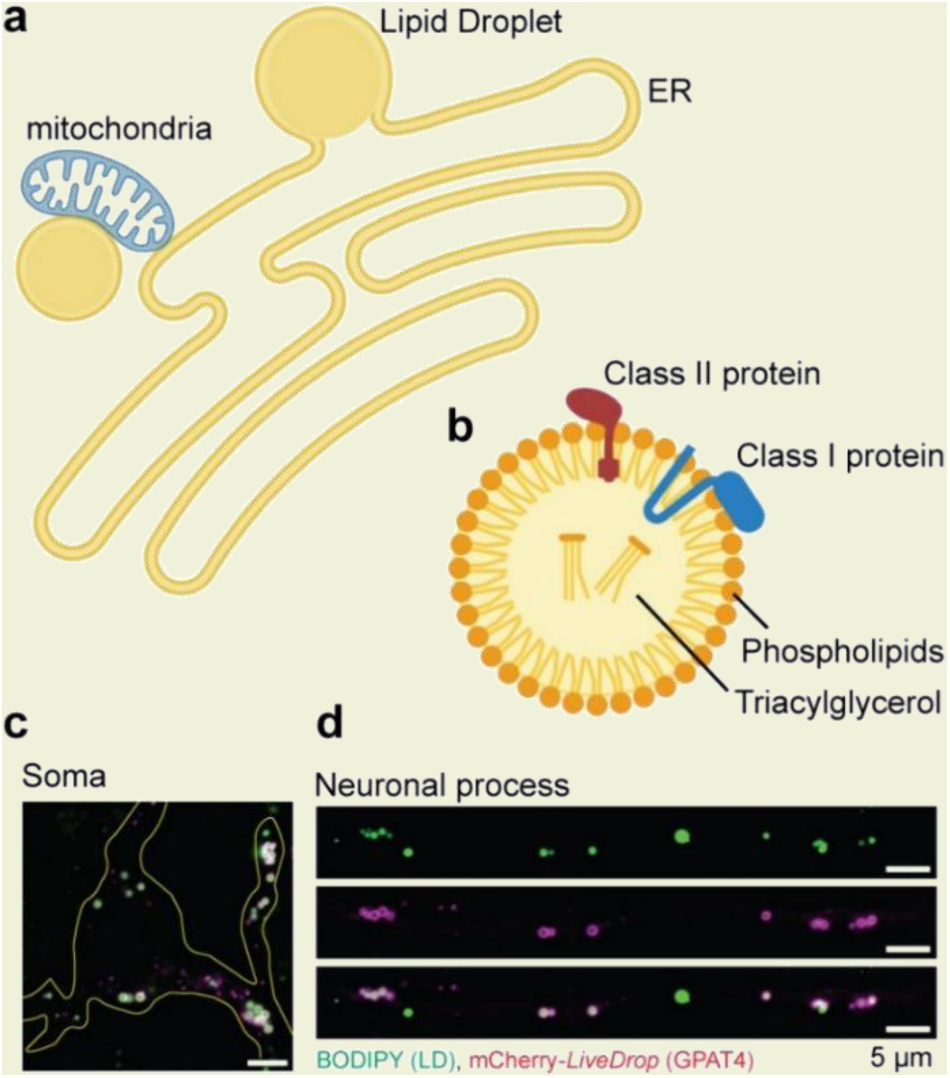

**a,** Schematic illustration of LD biogenesis at ER and its association with mitochondria. **b,** Architecture of LDs showing a TG-rich core surrounded by phospholipids and proteins. **c-d,** Fluorescence images of LDs in the soma (c) and neurites (d) labelled by BODIPY (green) and a model class-I LD protein mCherry-LiveDrop (GPAT4) (magenta).

Together, these observations underscore the need of a robust tool to interrogate NLDs with deep molecular resolution. Comprehending how LDs—and their associated proteins—drive neuronal dysfunction in NDs requires isolating LDs free of intracellular organelles such as mitochondria, ER, and lysosomes, with which they frequently interact. Here, we present a refined, easy-to-follow and highly reproducible protocol to facilitate the formation of mature LDs in cultured primary neurons and their isolation using a four-layer sucrose density gradient method. We also employed an LC-MS pipeline and developed analysis workflows to detect and quantify lipid species present in the core and monolayer of NLDs.

### Development of the Protocol

Lipid droplets as an organelle remained ignored for almost a century after their discovery in the 19^th^ century by Richard Altmann and E.B Wilson^31,32^. A series of discoveries in the 1990s outlined the importance of their role in cellular functioning and metabolic disorders, reigniting interest and prompted the development of innovative techniques for isolation of LDs from live tissues^2,33^. Irrespective of biological source, the core of LDs is mainly constituted of triglycerides and sterol esters that make them highly buoyant in aqueous medium, enabling their separation from other organelles by floatation. Initial attempts to systematically isolate LDs from plant-based sources date back to the 1970s^34,35^. Subsequent refinement of the methodology (Table 1) enabled extraction of the LDs in their purest form devoid of any membrane contamination from several animal sources for lipidomic and proteomic studies (Table 2).

**Table 1:**
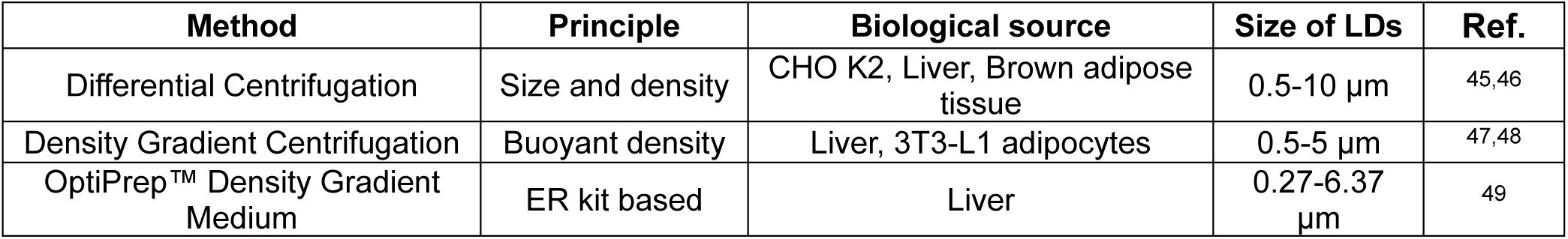
Methods for lipid droplet purification.

**Table 2:**
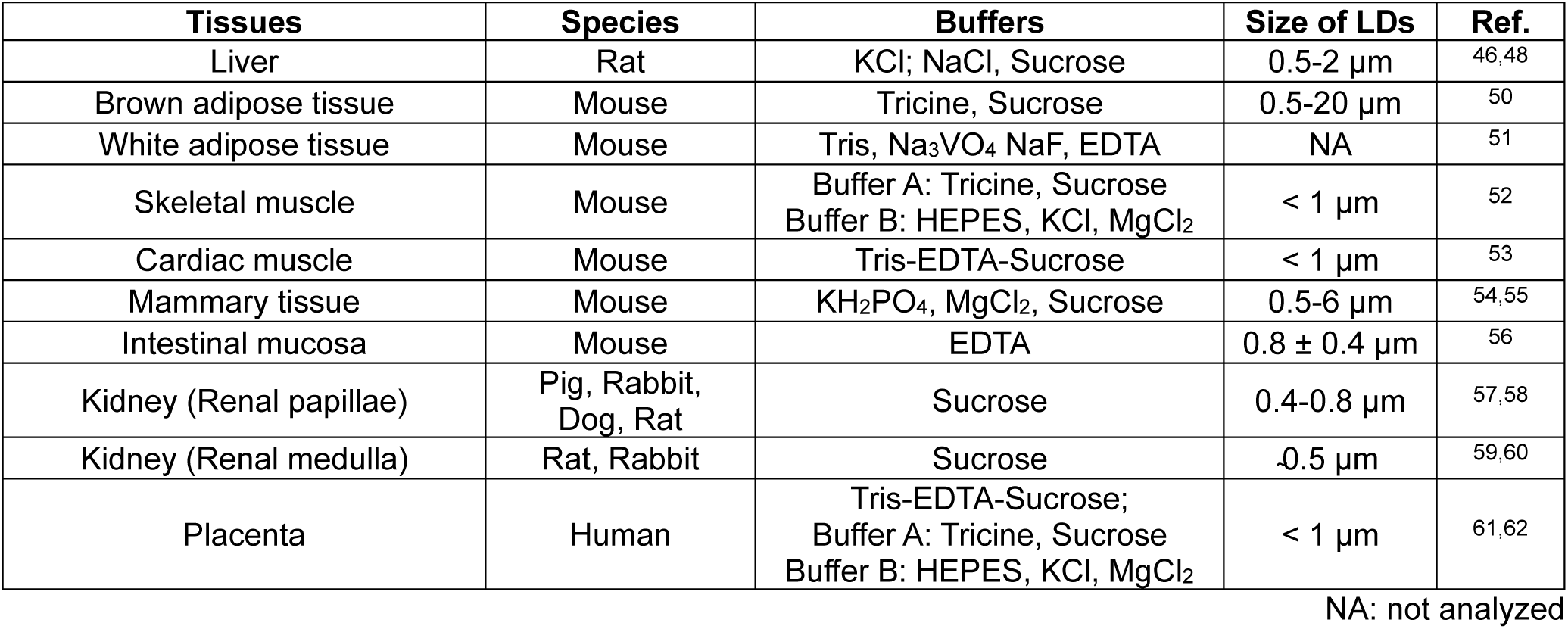
Purification of lipid droplets from animal tissues.

**Table 3:** Troubleshooting table.

| Step | Problem | Possible reasons | Potential Solutions |
| --- | --- | --- | --- |
| 5-8 | Underdeveloped neuronal processes | Neurons were not healthy | Plate neurons at adequate density and ensure the quality of culture media |
|  |  | Culture used earlier than DIV-14 | Allow neurons in culture for DIV-14 or more |
| 9 | Failure of LD induction | KLH45 was degraded | Reconstitute fresh KLH45, store at appropriate temperature in small aliquots and avoid freeze-thaw cycle |
| 12-14 | Incomplete lysis of neurons | Pressure of nitrogen in cell disruption vessel was low | Increase the pressure in nitrogen vessel |
| 23-28 | Low count of LDs | Incomplete lysis of cells | Ensure complete lysis of neurons by optimizing nitrogen cavitation in step 12-13 |
|  |  | Gradients layers of high volume | Use smaller tube for small samples |
|  |  | Loss during LD collection | Beckman OptiXTRACT Sample Recovery Aid can be used |
| 23-28 | Aggregation of LDs | Electrolyte in the solution was low | Check the concentration and pH of buffers |
|  |  |  | Additional floatation step in low concentration of glycerol, 0.1-1 M NaCl or 100 mM Na<sub>2</sub>CO<sub>3</sub> (pH 11.5)<sup>74</sup> |
| 28 | Disrupted LD membrane | Homogenization was too harsh | Reduce the nitrogen pressure |
| 28 | Contamination of membrane | Wide diameter tube used | Preferably use tubes of narrow diameter for prep. with small number of neurons |
|  |  | Density gradient was messy | Use smooth pipette with wide orifice tips on a sturdy platform |
| 36 | High phospholipids in TLC | Membrane contamination | Follow the above steps and ensure the quality of LDs before lipid extraction |
| 46 | Lack of two distinct layers or uneven layer volume across samples in MTBE:MeOH extraction of lipids | Improper preparation of the extraction solvent, for example, separation of MTBE:MeOH solvents prior to use | Ensure both the phase separation and extraction solvent are prepared correctly and properly mixed immediately prior to use |
|  |  | Sample buffer not compatible with extraction solvent | Test out a sample buffer that is primarily aqueous if necessary or use lower sample volumes, split and combine in drying step if necessary |
| 56-58 | Improper elution or poor retention time of lipids in TC | Leak in column attachments | Reattach column |
|  |  | LC-MS pump issue | Confirm that pressure gradient closely aligns with %B. As %B increases (more viscous solvents), the pressure should also increase. |
|  |  | Improper LC mobile phase preparation | Prepare fresh buffers if needed. |
| 60 | Retention time of a lipid species not consistent across samples. | Improper column heating | Check that column compartment is set to heat and is properly closed and maintained in a closed set. |
| 57-59 | Background signals in LC-MS that interfere with signal acquisition | Contamination of LC-MS solvents | If background signal is present in blank runs, the LC-MS solvents are likely contaminated. Replace LC-MS solvents. |
|  |  | Contamination within sample | If background signal is present in sample runs and extraction blank, contamination is likely occurring in extraction process. Check grade of all materials. Ensure that no plastic is used with strong acids and consider buffer-exchanging or washing the sample. |
| 59-60 | Lack of lipid identifications using lipid software | Improper MS acquisition parameters | Using MS vendor software, manually check the TIC to ensure there are peaks in MS and they are being selected for fragmentation |
|  |  | Poor fragmentation of lipid species | Collision energies and methodologies vary widely across mass spectrometers. Perform an energy titration or alternative fragmentation technique, if available on instrument. |

Isolation of pure LDs requires careful disruption of the cell membrane without rupturing internal membranous organelles. Moreover, the buoyancy of the LDs varies with the size of LDs (100 nm to more than 20 μm in some cell types) and their association with organelles (ER, mitochondria, endosomes etc.) prompted modifications in LD purification protocols to avoid co-floatation of undesirable membrane fragments. The present protocol was conceived to isolate NLDs that are physiologically representative, free of organelle contamination, and analytically compatible with LC-MS based Omics analysis. Different methods were employed in the past for the isolation of LDs from different tissues as outlined in Table 1, however complex morphology and low abundance of NLDs warranted significant improvements in the method for NLD isolation and quantitative estimation of the constituent lipids.

Earlier protocols described in Table 1 and 2 presented three persistent bottlenecks that could limit their application to neurons: (i) detergent-based lysis can disrupt LD-protein assemblies, whereas mechanical shearing may incompletely rupture neuronal processes and promote resealing of LDs within axonal membranes, (ii) LDs may also co-float with low-density organelles, often necessitating extensive washing with high-salt solutions that may degrade labile lipids, and proteins^33^, and (iii) the scarcity and inherently small size of NLDs lead to a low amount of lipid recovery, demanding high analytical sensitivity to achieve broad lipidome coverage in the LC-MS based analysis. To overcome these challenges, we integrated optimized nitrogen cavitation, a four-layer sucrose gradient in compatible buffers, and a global, untargeted LC-MS pipeline. Together, these refinements extended the scope of LD isolation from bulk tissue to primary neurons, enabling high-quality lipidomic characterization of NLDs.

We build upon a method previously developed for *in-vitro* motility assays that require intact lipid-protein interface of LDs^36^. *Barak et al.* optimized an LD isolation workflow with rodent liver and conducted optical-trap based motility assay that demonstrated the intact lipid-protein interface crucial for molecular motor driven transport of LDs. Subsequent refinements of the protocol by the same group demonstrated functional integrity of the LDs by showing that the isolated LDs, when bound to a microtubule, move in anterograde direction, an indirect proof of intact lipid-protein environment^36^. This protocol was further developed by *Rai et al.*, and *Kumar et al.* to show that the phospholipid composition of the hepatocytic lipid droplets changes during feeding to fasting transitions to regulate differential recruitment of kinesin motors^37,38^. These successive refinements established the utility of PIPES buffers for isolating LDs with preserved structure, enzymatic activity, and lipid integrity.

Protocols optimized for yeast and mammalian tissue samples report microgram range of proteins, corresponding to hundreds of micrograms of lipids given the 90:10 lipid-to-protein ratio in LDs. However, yield of NLDs is substantially lower, therefore we prioritized a high-yield and optimized lipid extraction built on methyl-tert-butyl-ether (MTBE) workflows^39^, high-yield resuspension solvents^40^, a robust LC- workflow, and specific data analysis programs^41,42^ (Fig. 5). We have further augmented the data analysis with an in-house program, written in Python. This enables a robust quantification and easy parsing of the data, both within a single lipid class and across the total sum of lipid classes. Such quantification of lipids, especially when looking at multiple classes, is critical because differences in ionization efficiency and adduct formation mean that the relative areas of two lipid species in the mass spectrum are not necessarily proportional to their relative concentrations. Internal standards are spiked in to account for losses during extraction and resuspension, and to correct for lipid class- and adduct-dependent differences in ionization efficiency. For each lipid species, the Python script assigns the internal standard with the closest retention time and identical adduct, thereby improving correction for ion-suppression effects. The known concentration of the standard and its area are then used to calculate the concentration of each lipid. Scripts and template files are all available on github at https://github.com/guptaMSlab/Lipidomics-Python-Quant-Protocol^43,44^. This approach delivers head-to-tail coverage of NLD lipids—revealing extensive species level characterizations.

### Overview of the Procedure

#### Step 1-9: Induction of LDs in cultured neurons

The workflow (Fig. 1) begins with dissection of the cortex from P0 neonatal mice pups, followed by dissociation and culture of primary neurons up to DIV14 under defined conditions, to allow maturation of neuronal processes. NLDs formation is then induced pharmacologically by KLH45, a cell-permeable small-molecule inhibitor of neuron enriched DDHD2-containing triglyceride lipase^12^. Successful induction of NLDs can be verified by staining neurons with neutral lipid dye, BODIPY493/503, followed by confocal microscopy and thin layer chromatography (TLC) (Fig. 2).

**Fig. 1:**
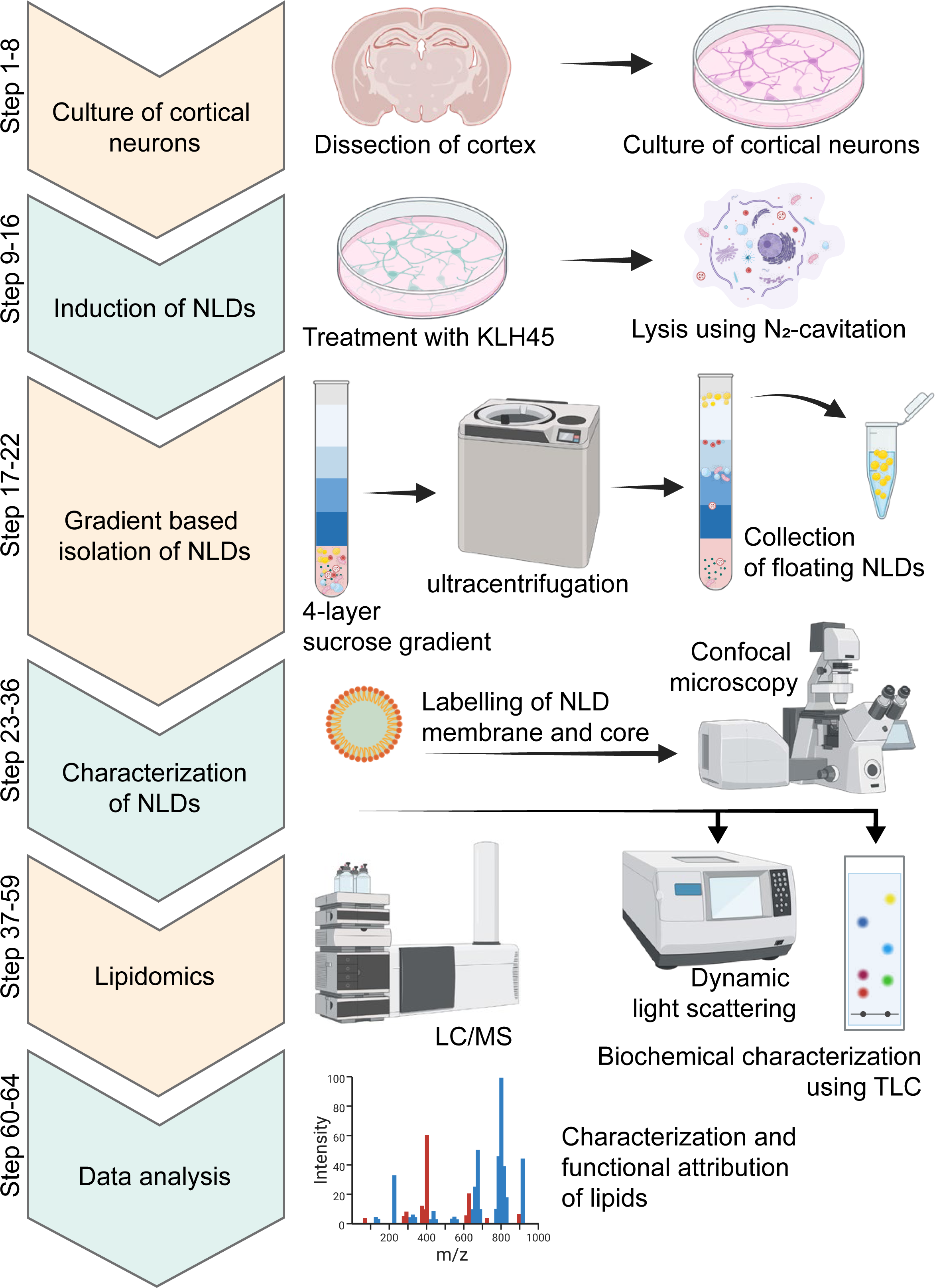
Overview of the procedure. Step 1-8: Dissect neonatal mouse cortex, dissociate, and culture primary cortical neurons to maturation. **Step 9-16:** Pharmacologically induce NLDs in cortical neurons and verify induction by staining with BODIPY493/503 (see figure 2). **Step 17-22:** Isolate buoyant NLDs by ultracentrifugation through a four-layer sucrose gradient and collect the floating fraction (see figure 3). **Step 23-36:** Characterize purified NLDs by confocal microscopy (labelling of neutral-lipid core and monolayer phospholipids using BODIPY493/503 and CellMASK respectively), dynamic light scattering (size distribution), and thin layer chromatography (class-level lipid quality control) (see figure 4). **Step 37-59:** Perform high-sensitive LC-MS with MS-compatible solvents to profile lipid species across classes (see figure 5). **Step 60-64:** Analyse spectra to identify and quantify lipid species using internal standards, generating class/group summaries and downstream statistical comparisons (see figure 6).

**Fig. 2:**
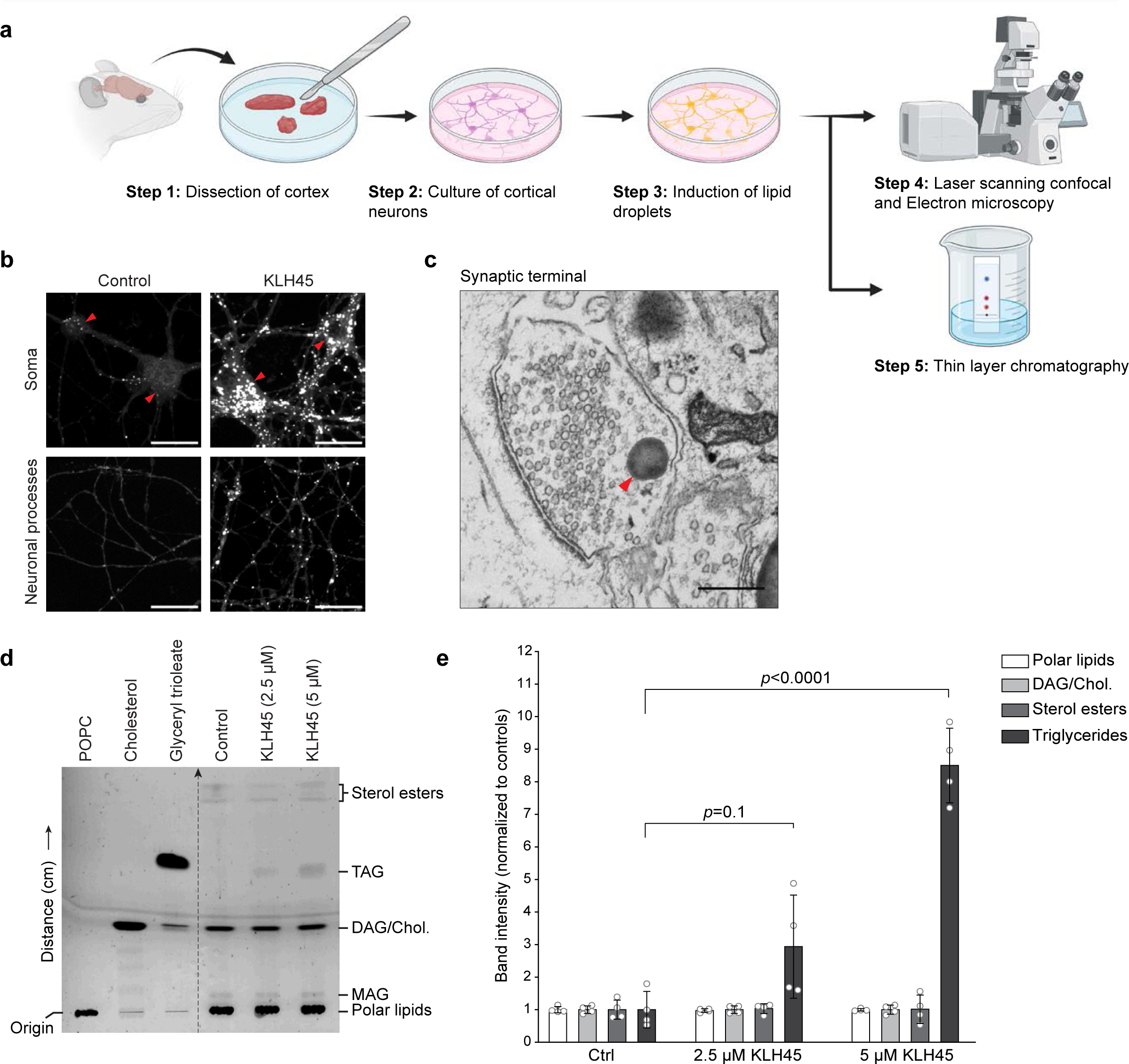
Induction of lipid droplets in dissociated cortical or hippocampal neurons. a, Schematic illustration of protocol steps to induce accumulation of LDs in the cultured primary neurons and confirmation by microscopy and thin layer chromatography (TLC). Cortical neurons from rodent brain cultured on poly-d-lysin coated dishes are treated with KLH45 at DIV 14-21. **b,** Confocal micrographs of control and KLH45 treated cortical neurons after BODIPY staining. Lipids droplets are apparent as globular structures both in the soma (red arrowheads) and neuronal processes after KLH45 induction. Scale bar: 20 µm. **c,** Electron micrograph of a hippocampal synaptic terminal from neurons treated with KLH45 (5 µM), showing LD accumulation (red arrowhead) positioned adjacent to a cluster of synaptic vesicles. This close apposition is consistent with the proposed role of LDs in local synaptic metabolism. Scale bar: 500 nm. **d,** TLC showing a dose-dependent accumulation of triglyceride in the cortical neurons after inhibition of neuron-specific triglyceride lipase with KLH45. Standard TG (glyceryl trioleate), cholesterol and phosphatidylcholine (POPC) are shown on the left side of grey line as references. **e,** Band intensity of lipid classes of control and KLH45 treated (2.5 µM and 5 µM) neurons are shown. Data from four independent experiments are represented as mean; error bars, s.d. *p*-values were determined using one way ANOVA with Tukey’s test. TG: triglyceride, DG: diglyceride, MG: monoglyceride, Chol.: cholesterol.

#### Step 10-22: Lysis of neurons and purification of LDs

Neurons are lysed by nitrogen cavitation, followed by slow pressure release to yield lysate that preserves the integrity of LDs. Nuclei and large cellular debris are cleared by table-top centrifugation to obtain a post-nuclear supernatant (PNS). To float the LDs, PNS is mixed with 2.5 M sucrose and overlaid with reducing density of sucrose in MEPS buffer. After ultracentrifugation, the floating NLD fraction is collected from the top, sucrose-free layer for physical characterization, and downstream biochemical analysis (Fig. 3).

**Fig. 3:**
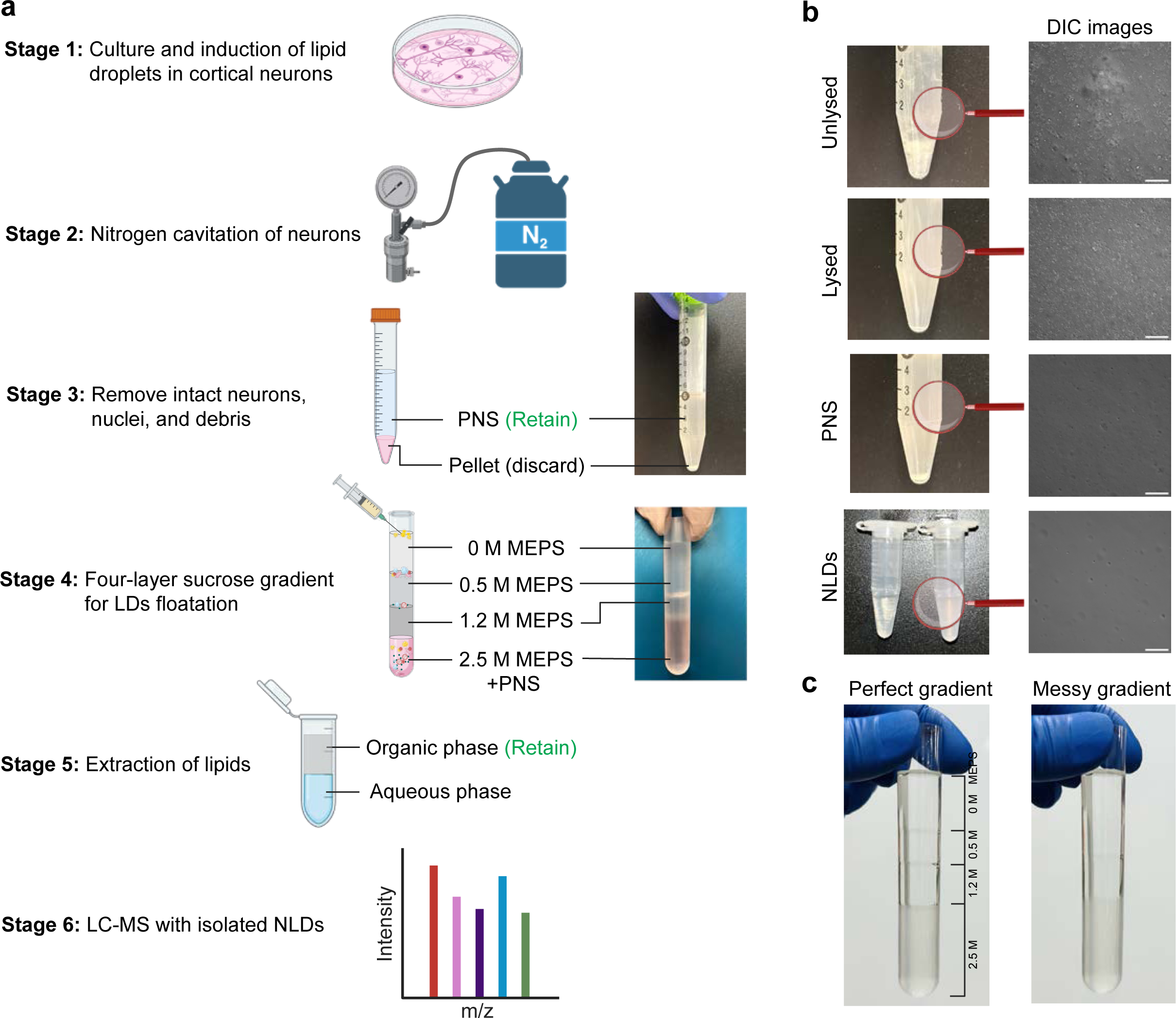
Workflow and key quality-control checkpoints for isolation of lipid droplets from cortical neurons. a, Schematic overview of the lipid droplet isolation workflow: **Stage 1,** LDs are induced in primary cultured cortical neurons. **Stage 2,** neurons are lysed by nitrogen cavitation. **Stage 3,** pellet containing intact cells, nuclei, and large debris are discarded after low-speed centrifugation, and supernatant is retained for the next step. **Stage 4,** NLDs are isolated by floatation through a four-layer sucrose gradient. The NLDs fraction is collected from the top layer (0 M MEPS) of sucrose density gradient. **Stage 5,** lipids are extracted, and the upper organic phase is retained. **Stage 6,** lipid composition is analyzed by LC-MS. **b,** Representative images of samples collected at successive stages of the workflow. Images on right side show high magnification views acquired using differential interference contrast (DIC) microscope. Scale bars, 50 µm. **c,** Examples of a correctly layered (perfect) and disturbed (messy) four-layer sucrose gradient. A properly layered gradient shows sharp interfaces between 2.5 M, 1.2 M, 0.5 M, and 0 M MEPS layers, whereas a disturbed gradient shows mixing between layers and should not be used for NLD purification.

#### Step 23-36: Quality control

Size distribution of the isolated LDs is determined by dynamic light scattering (DLS) method. The esterified lipid rich core and monolayer phospholipid of the LDs are stained by BODIPY493/503 and CellMask stains respectively, followed by fluorescent imaging on confocal microscope to confirm the integrity, and absence of membrane contamination in the floated NLDs. Two-step TLC, using organic solvents described in the chemical reagents section, is performed on isolated NLDs to confirm triglyceride enrichment and to survey other major lipid classes (Fig. 4).

**Fig. 4:**
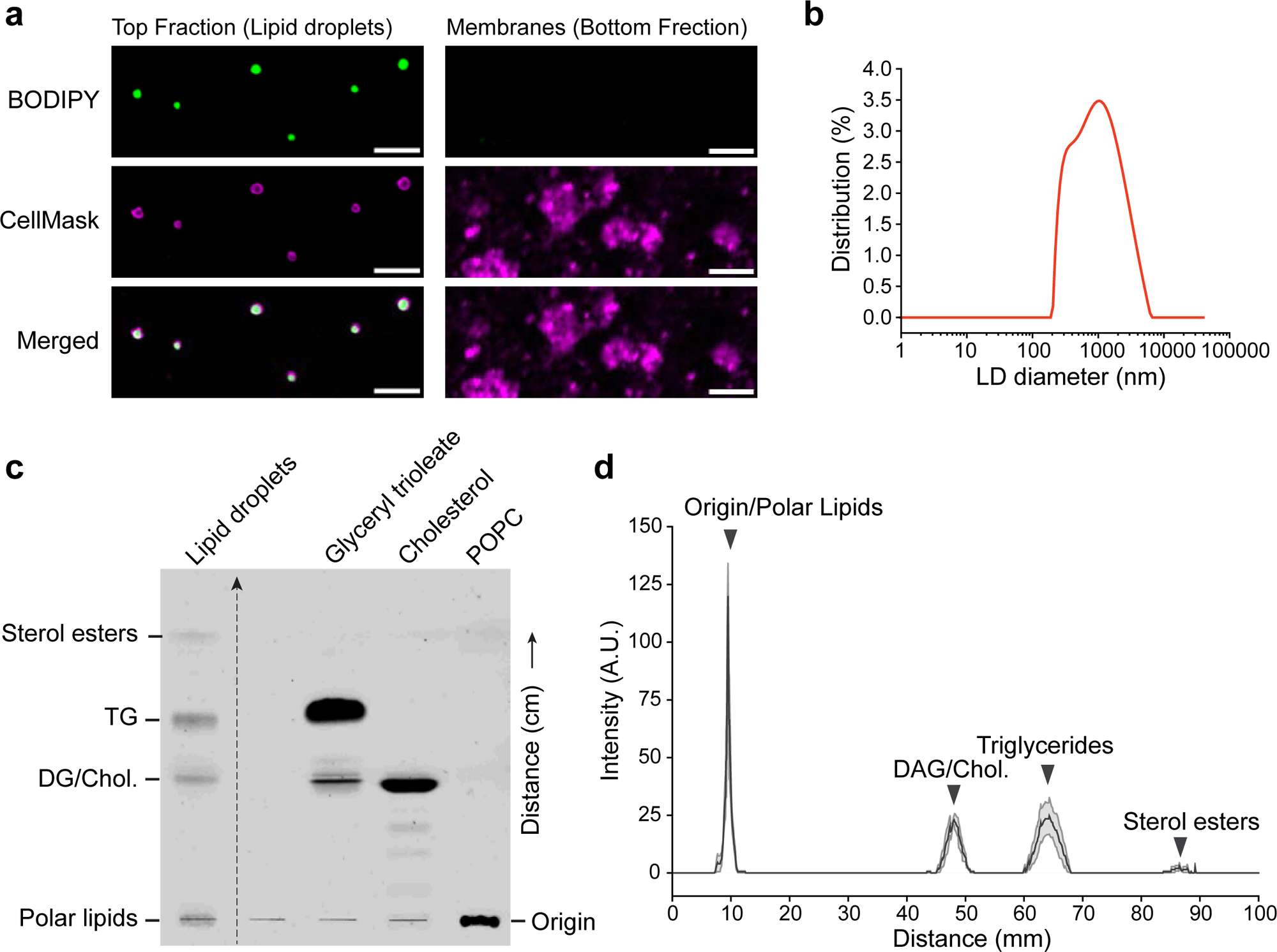
Physical and biochemical characterization of isolated neuronal lipid droplets. a, Representative confocal images of the top floating fraction (NLDs) and the bottom membrane-rich fraction after sucrose-gradient floatation. Samples were stained with BODIPY to label neutral-lipid core and CellMask to label phospholipid-rich membranes. Top fraction contains BODIPY-positive puncta outlined by CellMask, whereas bottom fraction contains only CellMask positive puncta without detectable BODIPY signal. Scale bars: 5 µm. **b,** Size distribution of NLDs analyzed by dynamic light scattering instrument (Anton Paar Lifesizer 500), showing diameters ranging between ∼200 nm to ∼5 µm. **c,** Thin layer chromatography (TLC) of lipids extracted from isolated NLDs. Major bands corresponding to sterol esters, TG, DG/chol., and polar lipids are indicated on the left side. Standard lipids were run on the right side for reference. **d,** Background-subtracted lane intensity profile of the TLC shown in c, demonstrating strong enrichment of triglyceride in the isolated NLDs, with lower signals corresponding to polar phospholipids, DG/chol. and sterol esters. The line-profile graph was generated from four independent experiments and is shown as mean; error bars, s.e.m.

**Fig. 5:**
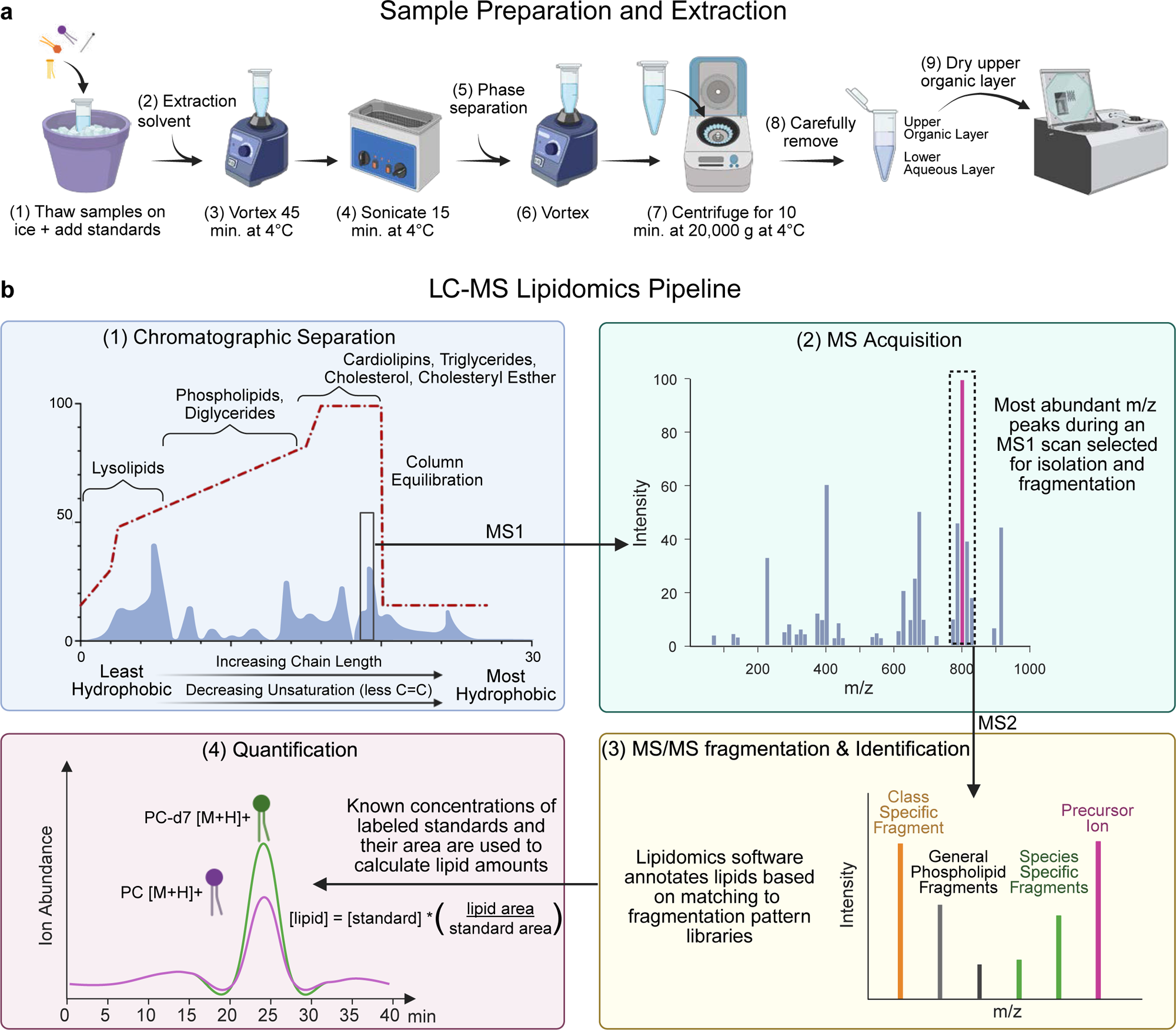
Schematic of MTBE lipid extraction and DDA-based LC-MS workflow. a, Schematic of the lipid extraction procedure from isolated NLDs, corresponding to **Procedure steps 37–48**. **Stage 1 (steps 37–38)**, thaw samples on ice and spike with internal lipid standards. **Stage 2 (step 39)**, add extraction solvent. **Stage 3 (step 40)**, vortex for 45 minutes at 4 °C. **Stage 4 (step 41)**, sonicate for 15 minutes at 4 °C. **Stage 5 (steps 42–43)**, add phase-separation solvent and mix thoroughly. **Stage 6 (steps 43–44)**, vortex and centrifuge to generate phase separation. **Stage 7 (steps 44–45)**, resolve the sample into an **upper organic layer** and **lower aqueous layer**. **Stage 8 (step 46)**, carefully collect the upper organic phase without disturbing the interface. **Stage 9 (steps 47–48)**, dry the upper organic phase and store the extracted lipids for downstream LC–MS analysis. **b,** LC-MS lipidomics analysis workflow, corresponding to procedure steps 49-64**. Stage 1 (steps 49–58)**, chromatographic separation: dried lipids are resuspended, loaded into LC-MS vials, and injected onto a C18 stationary-phase column for reverse-phase LC separation, in which an increasingly hydrophobic mobile phase elutes lipids according to their hydrophobicity. **Stage 2 (step 59)**, MS acquisition: the mass spectrometer, coupled online to the LC, acquires full-scan MS1 spectra across the specified m/z range and selects the most abundant precursor ions for fragmentation in a data-dependent acquisition (DDA) mode. **Stage 3 (steps 60–62)**, MS/MS fragmentation and identification: the resulting fragmentation spectra are matched against lipidomics libraries containing class- and species-specific precursor and fragment ions for lipid annotation. **Stage 4 (steps 63–64)**, quantification: identified lipids are quantified by comparing their peak areas with those of class-matched, isotope-labeled internal standards of known concentration.

#### Step 37-64: LC-MS Data acquisition and analysis

TLC predominantly detects neutral lipids as the major constituents of NLDs, whereas other lipid classes are not readily resolved under these conditions. To detect and quantify low abundance polar lipids, we developed a high-sensitivity data dependent analysis (DDA)-based LC-MS lipidomics workflow (Fig. 5). This approach combines a stringent lipid extraction protocol with streamlined acquisition workflows and a dedicated data-analysis program optimized for ease of quantification^63^. Resulting spectra are quality-filtered, and reference matched against curated lipid reference libraries for identification through class-based fragmentation, prior to quantification using spiked-in class and adduct specific internal standards (Fig. 5).

#### Applications of the method

Neurons are unique in their morphology and biochemical pathways. They harbor very few tiny LDs which in basal condition make the isolation and characterization challenging. To the best of our knowledge, this is first ever study where we successfully induced large LD buildup in dissociated neurons to purify them using a density-based floatation method. The lipidomic studies on purified NLDs provide a snapshot of the lipid milieu linked to neuronal LDs (Fig. 6).

**Fig. 6:**
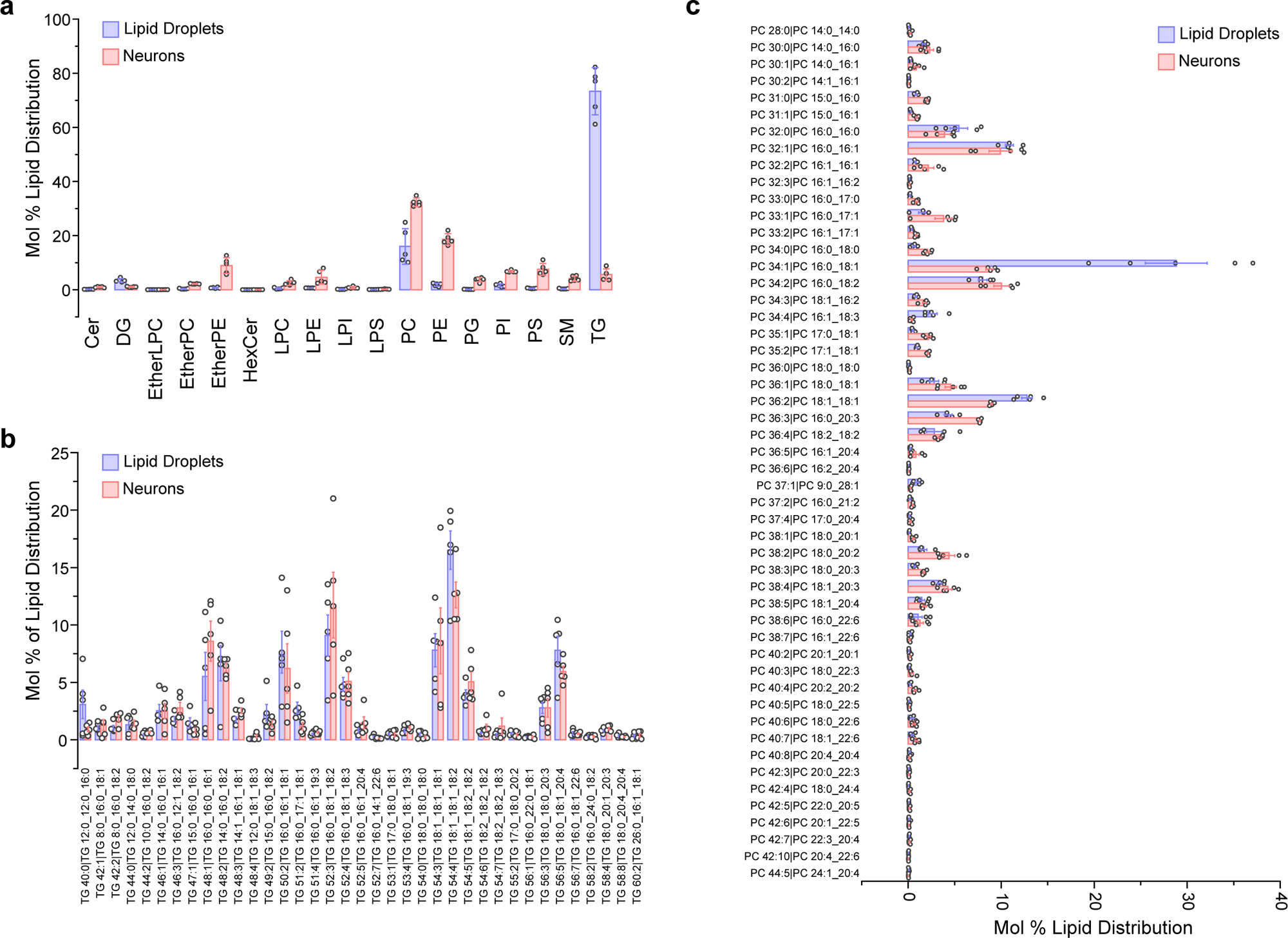
Lipidomic analysis of LDs purified from cortical neurons and whole neuron post-nuclear supernatant. a, Quantitative mol% distribution of lipid classes identified by LC-MS/MS from LDs and whole neuron. The lipid classes include Cer: Ceramides, DG: Diacylglycerol, EtherLPC: Ether-linked Lysophosphatidylcholine, EtherPC: Ether-linked phosphatidylcholine, EtherPE: Ether-linked phosphatidylethanolamine, HexCer: Hexosyl Ceramides, LPC: Lysophosphatidylcholine, LPE: Lysophosphatidylethanolamine, LPI: Lysophosphatidylinositol, LPS: Lysophosphatidylserine, PC: Phosphatidylcholine, PE: Phosphatidylethanolamine, PG: Phosphatidylglycerol, PI: Phosphatidylinositol, PS: Phosphatidylserine, SM: Sphingomyelin, TG: Triglyceride. The graph indicates mole % of total quantified lipid within the 5 independent samples represented as mean; error bar, s.e.m. In the lipid droplets, TG and PC comprise the major constituent lipids. **b-c,** Bar graphs representing the fatty-acyl species breakdowns among these two largest lipid classes (TG and PC respectively) and their distribution across neurons and lipid droplets. Data is represented as mean; error bar: s.e.m. Complete lipid profiles for all classes are attached as Source Data.

The step-by-step workflow described in this protocol provides an opportunity for neurobiologists to apply a diverse set of techniques to study lipid metabolism in neurons. Our successful integration of the method with high-throughput lipidomics provides an excellent avenue to identify targets of disease-specific alterations in the lipid, and in the future, protein composition of NLDs. The strategy of mass spectrometry-ready NLD purification will support high-throughput screening: small-molecule or genetic perturbations that can be ranked by their impact on NLD lipid classes, specific species, or surface-bound proteins. In conclusion, applications of the method described here can extend beyond simple cataloging to mechanistic dissection of how NLDs buffer the neuronal metabolic reserve. They can also be used to interrogate the NLD biogenesis and dynamics across neurodegenerative disease conditions, neuroinflammatory responses, and aging related lipid clustering in neurons.

In parallel, the NLD isolation strategies described can also be interfaced with orthogonal biochemical and bioanalytical approaches to understand the role LD in neuronal health and disease. First, comprehensive quantitative proteomics on purified NLDs with interactive correlation with lipids will provide a high-resolution snapshot of the NLD proteo-lipid milieu. Second, side-by-side comparisons across conditions (e.g., neuronal activity, metabolic stress, inflammatory response and diseased models) can pinpoint metabolic and disease-specific alterations in NLD composition and their binding partners, suggesting potential targets for corrective intervention. Third, combining the purified NLDs with assays of organelle crosstalk (e.g., LD-mitochondria, LD-ER, LD-lysosome interactions) and stable-isotope tracing (e.g. ^13^C-fatty acids) will enable flux-level insights into esterification, lipolysis, and acyl-chain remodeling in neurons.

#### Comparison with other methods

It is critical to highlight that the isolation of LDs from neurons presents unique difficulties and necessitated development of neuron specific protocols. Conventional methods for lipid droplet (LD) isolation, such as those described by *Ding et al.,* rely on mechanical disruption of cells^33^. While effective in many cell types, neurons present unique challenges due to their polarized structure with extending axonal processes away from the soma. Mechanical shearing ruptures the soma but only transiently perforates axons, which rapidly reseal, trapping LDs and vesicles within axonal membranes which would contaminate downstream processing. To overcome this limitation, we employed nitrogen cavitation, where rapid decompression of dissolved nitrogen produces isotropic microbubbles that uniformly rupture axons while minimizing shearing^64^.

Buffer composition also represents a critical determinant of LD integrity and analytical quality. Most of the previous protocols employed HEPES or Tris-based buffers (Table 2), which can chelate divalent ions, perturb membranes, and promote Na⁺/K⁺ adduct formation leading to background noise in mass spectrometry^65,66^. These issues necessitate additional desalting and washing steps that risk oxidative damage of lipids and sample loss—serious drawbacks when working with scarce NLDs. Therefore, we used PIPES-based buffers, which exhibit negligible metal binding and minimal adduct formation^67,68^. Previous methods including that by *Ding et al.* use single-layer sucrose cushions which may result in co-floatation with other LD interacting organelles, requiring additional washing steps^33^. Our adaptation of a four-layer sucrose gradient with defined salt concentrations allows LDs to ascend into a sucrose-free fraction while in-gradient washes strip off contaminating membranes. These approaches reduce isolation time, minimize sample loss, and yield clean NLDs that are directly compatible with downstream analyses.

#### Limitations of the method

Lipids on the monolayer membrane of LDs are synthesized at the cytoplasmic leaflet of the ER or on-site of maturing LDs. Proteins, on the other hand are trafficked from the ER membrane via membrane bridges or from the cytosol by amphipathic α-helices^30^ (Box 1). Certain lipids, and cytosolic proteins in particular, are susceptible to degradation or dissociation during the physical purification process leading to underrepresentation in downstream high-throughput studies. In addition, LD composition is size dependent^69^, and therefore analytical approaches that report population average may obscure size-resolved heterogeneity in neutral lipids, signaling lipids, and NLD-bound factors. Furthermore, because of low abundance of NLDs and their associated proteins, the current validation is based on imaging, flotation behavior, TLC, and lipidomic composition, rather than on purity defined by a conventional organelle protein-marker panel. Therefore, complementary biochemical and immunocytochemical methods in intact neurons are recommended to study the native localization, stoichiometry, and functions to assess size-dependent differences of NLDs.

Our isolation protocol typically requires large-scale primary neuronal cultures, limiting immediate application to scarce neuronal subtypes. Moreover, low-temperature handling—necessary to preserve lipids and reduce enzymatic turnover—can alter lipid-phase behavior and potentially shift the natural state of NLD lipid-protein complexes^70^. However, appropriate adaptations in the protocol can address these limitations without any impact on output. For example, immortalized neuronal lines or iNeurons can substitute for bulk primary cultures in pilot studies, while immunomagnetic capture of LDs using anti- perilipin2 (LD-coating protein) beads can enrich droplets even from limited samples. To control class-specific loss and extraction bias, samples can be spiked with stable-isotope-labeled lipid standards. Finally, to retain transient or weak protein interactors and enable correlative organelle studies, we recommend integrating proximity-label strategies with fluorescence-guided sorting of BODIPY-positive organelles. These adaptations will not just preserve biological fidelity but also improve quantitative confidence while acknowledging the intrinsic trade-offs of isolating a dynamic and biochemically heterogeneous cytosolic organelle, NLDs.

### Experimental Design

#### Neuronal model and culture scaling for downstream analysis

Low abundance of LDs in healthy neurons is a major experimental constraint: however, their abundance increases in altered metabolic conditions^12,24^. Therefore, the experimental design should be tailored to the biological source, LD abundance and downstream analytical objectives. Primary cortical or hippocampal neurons in step 1-8 should be maintained at least DIV 14 to ensure robust arborization of neurites and to enable the synapses to mature^71^. This maturation step is important because NLDs are closely linked to local glycerolipid metabolism within synaptic compartments^24,25^. For pilot experiments, iNeurons or immortalized neuronal lines may be used, but induction parameters in step 9 should be optimized for each model separately because basal LD abundance, responsiveness to lipase inhibition, and tolerance to lipid stress may vary significantly across neuronal subtypes. The scale of culture should be adjusted to match the intended downstream applications of the purified NLDs. Light microscopy-based validation of LD induction or electron microscopy in step 9 requires only modest input, whereas biochemical and lipidomic analyses at later stages require progressively larger preparations and stricter quality control. As a thumb rule, approximately 1X10^5^ neurons sparsely cultured on glass slides are sufficient for imaging experiments in step 9, 1X10^6^ neurons for TLC-based validation in step 29-36, and 10X10^6^ neurons for LC-MS experiments that aim to achieve broad lipidome and proteome coverage in step 37-59. Even smaller inputs can be used for the LC-MS, but it will necessitate more stringent lipid extraction and likely end up reduced depth of lipid detection.

#### Optimization of LD induction, neuronal lysis and LD floatation

Before starting actual experiment, three stages should be optimized in a pilot experiment: LD induction, lysis, and sucrose gradient described in step 9-22. KLH45 induces NLD accumulation in a dose-dependent manner (Fig. 2d,e); therefore, inhibitor concentration and treatment time should first be validated as in step 9 by BODIPY-staining before scaling up to large culture. Nitrogen cavitation in step 12 should be optimized for pressure and duration to achieve efficient lysis while minimizing disruption of intracellular organelles. Because organelle density and abundance may vary across the neuronal models, the ultracentrifuge tube capacity, volume of sucrose layers, and volume of top layer should be optimized during step 17-20. The isolated NLDs can be physically and biochemically characterized through light microscopy, dynamic light scattering, and TLC as described in steps 23-36 before downstream processing of the sample.

#### Lipid extraction and LC-MS analysis

NLDs yield small amounts of material; therefore, analytical reproducibility requires minimum losses in the process of lipid extraction during steps 37-48. To account for the contamination, losses, and adduct-dependent differences, deuterated internal lipid standards mixture (UltimateSPLASH^TM^ One) should be added to the sample in step 38. The amount and composition of internal standards should be chosen to match the expected class of lipids. All solvents used in step 37-48 should be prepared fresh and handled at low temperature to minimize oxidation and degradation of labile lipid species. For the low input NLD samples, excessive dilution at the resuspension step 50 can reduce the detection of minor lipids. At the same time, incomplete resuspension of the dried lipids can introduce quantitative variability. Moreover, vortexing, sonication, and clarification during steps 40-46 should be standardized across all samples. LC-MS acquisition parameters in step 59 should be optimized according to the instrument platform, detector type, ion optics, fragmentation mechanism, and sample complexity. High mass resolution improves the discrimination of lipid species with similar *m/z* values and enables more accurate lipid identification. Similarly, collision energy and the number of precursor ions selected during data-dependent acquisition (DDA) in step 59 should be optimized to balance fragmentation quality, lipidome coverage, and MS1 acquisition time. In this study, a top-3 DDA method with a fixed collision energy of 25 eV was selected to provide high-quality fragmentation spectra while maintaining robust MS1 acquisition for lipid quantification of the relatively less complex lipid droplet lipidome. Depending upon the instrument and the complexity of the sample, users may use different parameters.

#### Data processing and lipid identification

Accurate lipid identification depends on appropriate selection of the data-processing parameters according to the LC-MS platform, detector type, mass resolution, and fragmentation method. In this study, MS-DIAL was used for lipid identification, and a subset of LipidBlast *in silico* database was used for spectral matching during steps 60-62. Other suitable public or in-house databases may also be used in step 60. Following data processing, peak integration, fragmentation patterns, and retention time assignments should be reviewed to ensure accurate lipid identification prior to quantification in step 63- 64. For example, within a given lipid class (e.g., PC), longer-chain PC species with the same degree of unsaturation generally elute later, whereas more saturated PC species with the same chain length also elute later.

#### Control and replicates

Appropriate controls should be included throughout the workflow. Vehicle-treated or untreated neurons should be processed in parallel with induced samples to distinguish the treatment-dependent changes from basal lipid pools in step 9. An aliquot of post-nuclear supernatant should be retained at step 16 as an input control to evaluate enrichment and comparison with NLD lipidome. For statistically meaningful comparison, at least three biological replicates, each derived from independent neuronal preparation, are recommended. Moreover, technical repeats may be included during establishment of the method or instrument optimization.

#### Variations and adaptations

Although primary neurons were used during step 1-8 as the model system in present protocol, this modular workflow can be adapted according to sample availability and the biological question. In principle, this protocol can be applied to brain tissue samples with NLD accumulation. The main adaptation before LD floatation would require gentle dounce homogenization after washing at step 11 prior to nitrogen cavitation at step 12. For scarce neuronal populations or limited tissue samples, immunomagnetic enrichment of NLDs using anti-perilipin-2 beads can also be employed after step 15, but it would require additional standardization steps and specificity controls. Stable-isotope tracers or fluorescent lipids analogs can also be introduced before step 9 for the studies focusing on lipid flux rather than steady-state composition. In addition, proximity-labeling strategies or fluorescence-guided LD sorting strategies can be integrated before step 9 when the end goal is to define NLD-associated protein complexes or their interaction with other intracellular organelles.

#### Expertise needed to implement the Protocol

The protocol can be performed by researchers with experience in cell biology and biochemistry. Initial steps such as neuronal culture (steps 1-5), culture maintenance (step 6-8), LD induction (step 9), and imaging-based (step 9, 25-28) validation can be performed by users with basic training of neuronal culture and fluorescence microscopy. However, some stages such as nitrogen cavitation (step 12-13), setting up multi-layer sucrose gradient (step 17-19), recovery of floating LD fraction (step 21), and lipid extraction and TLC using organic solvents (29-36) require greater technical skills, practice, and caution. The LC-MS components of the protocol (step 37-64) require access to advanced instruments and should ideally be performed in collaboration with an experienced research team, or under the guidance of a researcher familiar with lipidomics workflow optimization and data analysis.

## MATERIALS

**BIOLOGICAL MATERIALS**

Animals: Wild-type strains of Sprague-Dawley rats (Charles River, Strain code 400, RRID: RGD_734476) and C57BL/6J mice (Jackson Laboratory, Strain code 000664, RRID: IMSR_JAX:000664) were used in our previous studies using this protocol^24^.

**▴CAUTION** All animal-related experiments were performed in accordance with protocols approved by the Weill Cornell Medicine and Yale University School of Medicine IACUC.

**REAGENTS**

- 1X PBS (ThermoFisher, cat. no. 10010031)
- Hanks’ Balanced Salt Solution, HBSS (ThermoFisher, cat. no. 14175095)
- MEM (Gibco, cat. no. 11095080)
- Fetal bovine serum, FBS (Gibco, cat. no. 16140071)
- D-glucose (Sigma-Aldrich, cat. no. G7021)
- Sodium pyruvate (Gibco, cat. no. 11360070)
- Neurobasal medium (Gibco, cat. no. 21103049)
- B-27 Supplement 50X (Gibco, cat. no. 17504044)
- L-glutamine (Gibco, cat. no. 25030081)
- GlutaMAX Supplement (Gibco, cat. no. 35050061)
- KLH45 (Sigma, cat. no. SML1998)
- PIPES (Sigma, cat. no. P6757)
- EGTA (Sigma, cat. no. E3889)
- MgSO_4_ (Sigma, cat. no. M3409)
- KOH (Sigma, cat. no. 221473)
- Complete Protease Inhibitor Cocktail (Roche, cat. no. 45-11697498001) **▲CRITICAL** Prepare fresh before use.
- PMSF (Sigma, cat. no. P7626) **▲CAUTION** PMSF is acutely toxic. Use disposable gloves while handling.
- Sucrose (Sigma, cat. no. S0389)
- Glyceryl trioleate (Sigma, cat. no. T7140)
- CellMask^TM^ plasma membrane stain (Invitrogen, cat. no. C10045)
- BODIPY493/503 (Thermo Scientific, cat. no. D3922)
- CuSO_4_ (Supelco, cat. no. 102791)
- H_3_PO_4_ (Sigma, cat. no. 345245)
- Methyl-tert-butyl-ether (MTBE) (Sigma, cat. no. 306975)
- Isopropanol, LC-MS grade (IPA) (Fisher, cat. no. A461-4)
- Acetonitrile, LC-MS grade (ACN) (Fisher, cat. no. A955-4)
- Methanol, LC-MS grade (MeOH) (Birch, cat. no. 1935-5)
- Butanol, HPLC grade (Thermo Scientific, cat. no. 71-36-3)
- Formic Acid, LC-MS grade (Thermo Scientific, cat. no. T85178-AD)
- Ammonium Acetate (MP Biomedicals, cat. no. 198759)
- Organic solvents: Chloroform (Sigma, cat. no. 288306); Methanol (Sigma, cat. no. 34860); n-Hexane (Sigma, cat. no. 34859); Diethyl ether (Sigma, cat. no. 309966); Acetic acid (Sigma, cat. no. 320099) **▲CAUTION** The organic solvents are highly toxic and unstable explosive. Avoid contact with flames and always handle them in the fume hood. Dispose them in accordance with the institutional and local guidelines.
- Lipids: Glyceryl trioleate (Sigma, cat. no. T7140); UltimateSPLASH^TM^ ONE Lipidomics Mass Spec standard (Avanti Research, cat. no. 330820) **▲CRITICAL** Lipids are unstable and prone to oxidation under atmospheric conditions, so lipids should be purged with N_2_ gas before the storage. Avoid using plastic or polymer tubes for storage of the lipids. Use glass vials for the storage of lipids dissolved in chloroform or any other organic solvent.

**PLASMID**

- hSyn-mCherry-LiveDrop (GPAT4): Plasmid (Addgene plasmid number 250675, RRID: Addgene_250675) designed for this study.

**EQUIPMENT**

- Cell disruption nitrogen chamber (Parr Instrument, cat. no. 4639)
- Ultracentrifuge (Beckman Coulter, Optima L-100 XP ultracentrifuge)
- SW40 Ti rotor (Beckman Coulter, cat. no. 331302)
- SW40 14-ml ultracentrifuge tube (Beckman Coulter, cat. no. 331374)
- OptiXTRACT Sample Recovery Aid (Beckman Coulter, cat. no. D17685)
- Vortex Mixer (Scientific Industries, cat. No. SI-0236)
- Ultrasonic Bath (Branson, cat. no. CPX-952-217R)
- Vacuum Centrifuge (Savant, Speedvac Concentrator. Model NO. SVC-100H)
- Particle analyzer: Anton Paar Litesizer 500 equipped with Kalliope 2.8.0 software
- Microscope: Zeiss LSM880 laser scanning confocal microscope
- Agilent Poroshell 120 (EC-C18, 2.1 x 100 mm, 2.7 μm, 1000 bar) HPLC column
- Agilent 1290 Infiniti II LC system
- Agilent 6546 LC/Q-TOF mass spectrometer

**CONSUMABLES**

- Needles and plastic syringes (BD Biosciences)
- Hamilton syringes: 1-10 μl (cat. no. 80360), 10-100 μl (cat. no. 81085), and 0.1-1 ml (cat. no. 81330)
- Microcapillary pipets: Kimble Glass (cat. no. 71900-25)
- TLC plates (Millipore, cat. no. 1.05721.0001)
- LC-MS vials (Thermo Scientific, cat. no. 6PMCK30LVW)

**SOFTWARES**

- MS-DIAL, version 5.3 (RRID: SCR_023076)
- ImageJ software, version 1.54f, https://imagej.nih.gov/ij/ (RRID: SCR_003070)
- Zen microscopy software, version 3.1 (RRID: SCR_013672)
- Python programming language, version 3.1 (RRID: SCR_008394)
- https://github.com/guptaMSlab/Lipidomics-Python-Quant-Protocol</objidref>
- Pandas, version 2.2 (RRID: SCR_018214)
- Kalliope 2.8.0 software

**REAGENT SETUP**

### [H3] Poly-d-lysins

Dissolve poly-d-lysine in sterile water to a final concentration of 50 μg/ml and sterilize the solution using 0.22 μm filter. The solution can be stored at 4 °C for up to 1 month. Warm the solution to room temperature (25 °C) before use; discard if there is any turbidity or contamination.

### Neuronal plating medium

Prepare neuronal plating medium by supplementing Minimum Essential Medium (MEM) with 10% (vol/vol) heat-inactivated FBS, 30 mM D-glucose, and 1 mM sodium pyruvate. Sterile-filter the medium if necessary and store at 4 °C for up to 1 month. Warm the medium to 37 °C before use.

### Neuronal maintenance medium

Prepare neuronal maintenance medium by supplementing Neurobasal Medium with 2% (vol/vol) B-27 and 0.5 mM L-glutamine. Alternatively, 0.5 mM GlutaMAX Supplement may be used in place of L-glutamine to improve glutamine stability during long-term neuronal culture. Prepare the medium under sterile conditions and store at 4 °C for up to 2 weeks. Warm the medium to 37 °C immediately before use.

### KLH45

Add 2.18 ml dimethyl sulfoxide (DMSO) to a vial of 5 mg KLH45 to prepare a 5 mM stock solution. Vortex for 5 min, aliquot in small sizes, and store at -20 °C for up to 6 months. Once thawed, aliquots should not be returned to storage for reuse.

### MEPS buffers

Prepare 0.9 M, 2.5 M, 1.2 M, 0.5 M & 0 M sucrose solutions (M denotes molarity of sucrose; 0 M MEPS indicates MEPS buffer without sucrose) in ultrapure water. Add PIPES (35 mM, pH 7.2), EGTA (5 mM) and MgSO_4_ (5 mM) to the sucrose solutions and maintain the pH to 7.2. The filtered stocks solutions can be stored at 4 °C for up to 6 months after preparation. Add freshly prepared protease inhibitor cocktail (1X) immediately before starting the experiment.

**▲CRITICAL** PIPES is not readily soluble in water at pH ≤ 7. Prepare a 1 M stock solution of PIPES by dissolving it in water and, while stirring, adjust the pH to 7.2 using KOH pellets. To facilitate the solubilization of 2.5 M sucrose, immerse the tube in a water-bath set at temperature 70 °C. Sucrose solutions can be stored for up to 6 months at 4 °C. Discard the solutions if any turbidity or contamination is observed.

### TLC solvent systems

Prepare 208 ml TLC ‘solvent A’ by mixing 120 ml n-hexane, 80 ml diethyl ether and 8 ml acetic acid (ratio 60:40:4); and 240 ml TLC ‘solvent B’ by mixing 236 ml n-hexane and 4 ml diethyl ether (ratio 59:1).

**▴CRITICAL** TLC ‘solvent A’ and ‘solvent B’ should be prepared and stored at room temperature (25°C) in narrow-mouth glass containers tightly sealed with glass stopper. The solvents can be used for up to 6 months unless exposed to air that may change the composition of solvents.

### Extraction solvent

Add 5 ml methanol to 15 ml methyl tert-butyl ether (MTBE) to prepare 20 ml extraction solvent.

**▲CRITICAL** The solvent should be prepared fresh before use. It is highly volatile; therefore, headspace in the storage bottles should be minimized, and the bottles should not be left open to air.

### Phase separation solvent

Add 15 ml ultrapure water to 5 ml methanol to prepare 20 ml phase separation solvent.

**▲CRITICAL** The solvent should be prepared fresh before use. As changes in solvent composition can compromise phase separation and reproducibility during lipid extraction, mix the solvent before use.

### Mobile phase A

Prepare required volume of ‘mobile phase A’ by mixing LCMS-grade acetonitrile:milliQ water (60:40, vol/vol) and supplementing the final mixture with 0.1% (vol/vol) formic acid and 7.5 mM ammonium acetate. It should be prepared at room temperature for immediate use.

### Mobile phase B

Prepare required volume of ‘mobile phase B’ by mixing LCMS-grade Isopropanol:LCMS-grade acetonitrile (90:10, vol/vol) and supplementing the final mixture with 0.1% (vol/vol) formic acid and 7.5 mM ammonium acetate. It should be prepared at room temperature for immediate use.

**PROCEDURE**

**Induction of LDs in cultured neurons**

**Primary culture of Cortical neurons**^72,73^ **●TIMING 15 d (6 h hands-on time)**

1. Coat 10 culture dishes of 150 mm size each with 10 ml of 50 μg/ml poly-d-lysine for 16 hours before starting the procedure of cortical culture.

2. Decapitate postnatal day 0 (P0) mouse or rat pups, dissect both the cerebral cortices, and immediately transfer the tissue to ice cold HBSS.

**▲CRITICAL STEP** Dissect out and remove the parts of the brain other than cortex to ensure the purity of the cortical neurons. Other regions of the brain such as hippocampus, cerebellum etc. can also be cultured separately using specific protocols for comparative study.

3. Slice the cortices into small pieces (∼1 mm) using a dissection knife and allow digestion in HBSS solution containing papain (20 U/ml) and DNase (20 μg/ml) for 15-20 minutes at 37 °C.

4. Triturate the digested tissue slices by pipetting back and forth with 1 ml tip and filter the debris out using a 40 μm cell strainer.

**▲CRITICAL STEP** Cell strainer with wider mesh size may allow cell debris to pass through, while ones with narrower mesh size will restrict the passage of dissociated cortical cells. Wet the strainer with 5 ml HBSS before pouring cell suspension to prevent clogging.

5. Wash the dissociated cortical cells with HBSS and plate them onto the poly-d-lysine coated petri dishes with 20 ml neuronal plating medium at 12,000-20,000 cells/cm^2^ density.

6. Exchange the plating media after 3 hours with 25 ml neuronal maintenance media containing B-27 and L-glutamine.

7. Maintain the cortical neurons at 37 °C in humidified incubator containing 5% CO_2_.

8. Replenish 30% of the total volume with prewarmed neuronal maintenance media at DIV4, 7 and 14.

**Induction of LDs in cortical neurons ●TIMING 1 d (30 min hands-on time)**

9. Add KLH45 to cortical neurons cultured in neuronal maintenance media to a final concentration of 5 μM at DIV14–DIV21, then return the culture dishes to incubator for 24 hours.

**▲CRITICAL STEP** To confirm the activity of KLH45 and successful induction of NLDs, stain untreated control and KLH45-treated neurons cultured on chamber slides with BODIPY 493/503 (follow step 25) and image using fluorescence or confocal microscope (Fig. 2b). For higher resolution, electron microscopy can also be performed (Fig. 2c). For detailed EM protocol, see also our published report ^24^.

**Lysis of neurons and purification of LDs**

**Disruption of cortical neurons ●TIMING 1.5 h**

10. Rinse the neurons twice with 10 ml ice cooled 1X PBS.

11. Add 400 μl 0.9 M MEPS buffer (ice cooled, supplemented with 1X protease inhibitor cocktail and 0.2 mM PMSF) to the layer of cortical neurons and scrape using a sterile plastic cell-scraper.

12. Transfer the cells to a pre-cooled cell disruption vessel. Dissolve oxygen-free nitrogen gas to the cytosol for 20 minutes under high pressure (750 pounds per square inch, psi).

13. Connect the release port to a collection chamber on ice by silicon pipe and release the pressure slowly to collect the disrupted neurons.

14. (*Optional*) Mount 30 μl lysed neurons between two glass slides (thickness: 0.18 mm) and observe on DIC microscope at 40X magnification to confirm the lysis of cortical neurons (Fig. 3b).

**▲CRITICAL STEP** Higher pressure may lead to complete homogenization of the cells as well as membranous organelles, therefore optimization of the pressure and time for N_2_ cavitation is required to achieve near complete lysis of neurons.

**Separation of intact neurons, ruptured cell membrane and nucleus ●TIMING 15 min**

15. Transfer the lysed neurons into a fresh 15 ml falcon tube and centrifuge at 1800 g for 10 minutes at 4 °C.

16. Transfer the supernatant (called post-nuclear supernatant or PNS, Fig. 3a-b) to a fresh tube and place it on ice while preparing the sucrose gradient. Optionally, retain an aliquot of 500 μl PNS on ice, or snap-freeze in liquid nitrogen and store at -80 °C for subsequent assessment of purification efficiency and comparison with isolated NLDs during lipidomic analyses.

**Purification of LDs by density-based floatation ●TIMING 3 h (1 h hands-on time)**

1. Transfer 3 ml 2.5 M MEPS to the bottom of a clear SW-41 centrifuge tube and overlay 2-3 ml PNS over it. Gently mix the two layers with a cut pipette tip to avoid frothing.

**▴CRITICAL STEP** Mix MEPS (2.5 M) and PNS (prepared in 0.9 M MEPS) in a ration that the final concentration of sucrose in the bottom layer is ≥ 1.5. Divide the PNS equally between two SW-41 tubes that can be easily balanced against each other during ultracentrifugation.

18. Overlay the bottom layer with 2 ml each of 1.2 M and 0.5 M MEPS from the bottom to top carefully without disturbing the interfaces.

**▲CRITICAL STEP** Use a smooth pipette with cut-tips to layer the sucrose gradients. A messy layering of the sucrose solutions will result in the incomplete isolation of LDs and contamination of other organelles (Fig. 3c).

19. Overlay 2-3 ml of 0 M MEPS over the layer of 0.5 M MEPS.

**▲CRITICAL STEP** The volume of the top layer of 0 M MEPS should be up to the maximum permissible height of the SW41 tube. Increasing the volume of the top layer improves the purity of the isolated NLDs by increasing the flotation distance. Optionally, salts like 100 mM sodium carbonate can be added to the top layer to remove the loosely bound non-specific proteins from the LDs^74^.

20. Transfer the SW-41 tubes to pre-cooled (at 4°C) SW-41-Ti swinging buckets and centrifuge at 174,000 g for 1.5 hours at 4 °C.

21. Transfer the buckets onto ice and collect the floating top layer of LDs using 25-gauge slanted needle attached to a 1 ml syringe.

**▲CRITICAL STEP** Collect the LD sample with minimum possible volume of 0 M MEPS buffer without disturbing the underlaying layers containing undesired membranes and cytosolic proteins. If the LDs appear dilute, centrifuge the sample at 14,000 g at 4 °C for 10 minutes and carefully remove the buffer at the bottom using a fine needle without disturbing the floating LD layer on the top.

22. Snap freeze aliquots of the isolated LDs samples in liquid nitrogen and store at -80 °C to preserve the lipids and LD associated proteins until further use.

**▪ PAUSE POINT** The LDs samples can be used within 6 months after preparation. Always thaw the LD samples on the ice before use.

**Quality control**

**Dynamic light scattering ●TIMING 1 h**

23. Dilute the LD samples in ultrapure water (1:50) and load onto dynamic light scatter particle analyzer (Anton Paar Litesizer 500).

24. Use the Kalliope 2.8.0 software to acquire the size distribution of LDs following manufacturer’s guidelines (Fig. 4a).

**Confocal micrography ●TIMING 4 h (1 h hands-on time)**

25. Prepare a staining solution containing 0.5X CellMask™ PM Stain (from 1000X stock) and 1X BODIPY 493/503 (from 1 mg/ml, i.e. 1000X stock) in 1X PBS.

26. Transfer 10 μl of LDs sample or 1:100 diluted membrane samples collected from the interface of lower sucrose gradients to two separate tubes containing freshly prepared staining solutions and incubate for 10 minutes at room temperature (∼25 °C).

27. Mount the stained samples on clean glass slides under coverslips and seal the edges with clear nail polish.

**▴CRITICAL STEP** Washing of the unbound CellMask™ and BODIPY493/503 stains is not required. The optimum concentration of the stains may vary between 0.5X to 2X depending on the size and density of the LDs.

28. Image the samples on confocal microscope equipped with appropriate lasers for CellMask™ and BODIPY493/503 stains and process the images using ImageJ software (Fig. 4b).

**Thin layer chromatography ●TIMING 2 d (6 h hands-on time)**

29. Thaw an aliquot (100-150 μl) of purified LD sample on ice, adjust the volume to 500 μl with ultrapure water and transfer it to a clean glass tube.

30. Add 2 ml methanol and 1 ml chloroform to the tube, vortex the mixture for 1 minute and place it at 4 °C for 14-16 hours.

31. Add 1 ml each of chloroform and water to the tube, vortex it again for 1 minute and allow it to sit undisturbed at room temperature for 30 minutes, or until the lower lipid-containing organic phase is distinctly separated from upper protein-containing aqueous phase.

**▲CRITICAL STEP** If the protein content of the sample is too high, the phases will not separate well under gravity. To separate such samples, prepare the sample in a glass centrifuge tube and spin at 100-200 g for 10 minutes at 4 °C.

32. Transfer the organic phase to another clean glass tube and evaporate the solvent under a stream of oxygen-free nitrogen gas.

33. Resuspend the dried lipids in 30 μl chloroform and spot them onto a silica TLC plate that has been pre-cleaned with chloroform.

34. Separate the spotted lipids by two-solvent systems A and B. First, develop the plate halfway in solvent system A [n-hexane:diethyl ether:acetic acid (60:40:4, vol/vol)], remove the plate, and allow the solvent to completely evaporate in the chemical fume hood. Next, develop the plate in solvent system B [n-hexane:diethyl ether (59:1, vol/vol)] to the top of the plate. Remove the plate and allow the solvents to evaporate completely again from TLC plate in the chemical hood.

**▲CAUTION** The solvent systems contain highly flammable, volatile, and corrosive chemicals. Handle and dry TLC plates in a fume hood, away from heat, sparks and open flames. Ensure complete evaporation of solvents before proceeding to the next step.

**▪ PAUSE POINT** The plates can be stored in a dry place for 7-10 days.

35. Visualize the lipids by spraying the TLC plate with a solution of 10% CuSO_4_ and 8% H_3_PO_4_ followed by charring the plate for 15-20 minutes in a preheated oven at 180 °C (Fig. 4c).

**▲CRITICAL STEP** To monitor specific lipids in the sample, spot standard lipids on separate spots on the level of other samples on the same plate. In case of any suspicion of lipid contamination in solvents, include a spot of solvent blank.

36. Capture the image of the TLC plate using a grayscale scanner and quantify the band intensities along the lipid migration lengths using ImageJ software (Fig. 4d).

**LC-MS Data acquisition and analysis**

**Extraction of lipids for LC-MS ●TIMING 1 d (1.5 h hands-on time)**

37. Thaw LD samples on ice and transfer to 2 ml click-seal microcentrifuge tube.

**▲CRITICAL STEP** Keep sample volume ≤ 200 μl to ensure complete sealing of the tubes. Prepare an ‘Extraction Blank’ (buffer and standards only, but no sample).

38. Spike each sample with 5 µl of internal lipid standards (i.e. UltimateSPLASH^TM^ One) before extraction. This corresponds to 40:1 (v/v) sample to internal standard for sample volumes 200 µl.

**▴CAUTION** UltimateSPLASH^TM^ One is supplied in volatile solvent (1:1 dichloromethane:Methanol) that may drip or readily dry up. Therefore, optimal care should be taken to prevent any loss while adding the standard to the samples.

**▲CRITICAL STEP** The volume of the internal standard can be adjusted to ensure that the signal remains within the dynamic range of LC-MS instruments.

39. Add 1 ml ice-cold Extraction Solvent [MTBE (methyl-tert-butyl ether): MeOH (3:1, vol/vol)] to each tube and tighten the caps immediately.

**▲CRITICAL STEP** Ensure the extraction solvents is properly mixed prior to use as the components can separate over time.

40. Vortex 45 minutes in cold room.

**▲CRITICAL STEP** Maintain cold conditions to minimize oxidation of lipids.

41. Sonicate in ice cold ultrasonic water bath for 15 minutes.

42. Add 650 μl phase separation solvent [H_2_O: MeOH (3:1, vol/vol)] to each tube and tighten the caps quickly.

**▲CRITICAL STEP** Ensure phase separation solvents are properly mixed prior to use as the components can separate over time.

43. Vortex for 2 minutes to integrate phase separation solvents.

44. Place into pre-cooled centrifuge and centrifuge at 20,000 g for 5 minutes at 4°C.

45. Remove tubes gently from centrifuge without disturbing layers.

**▲CRITICAL STEP** Visibly confirm the presence of two distinct layers – a lower aqueous layer and an upper organic layer. Some metabolites may partition at the interface between the two layers, leading to a thicker band between the two layers.

46. Gently pipette the top layer without disturbing middle or lower layer, and transfer the upper layer into a separate, fresh, centrifuge tube.

**▲CRITICAL STEP** Do not aspirate the intermediate layer as it contains salts and proteins that will subsequently suppress ionization. To improve comparability, collect same volume of the top fraction for all replicate samples.

47. Dry the collected organic phase at room temperature in a centrifugal vacuum concentrator compatible with organic solvents. Depending on sample concentration and vacuum efficacy, it may take 1-2 hours to overnight.

48. Purge the tube briefly with nitrogen gas, cap with parafilm, and store at -20°C in a box to protect from light exposure.

**▪ PAUSE POINT** Dried lipids can be stored at -20°C for up to a month.

**DDA based LC-MS data acquisition ●TIMING 30 min per sample**

49. Prepare resuspension solvent [1-butanol:IPA:H_2_O (8:23:69, vol/vol)]^75,76^. The solvent can be stored for ≤1 week at 4°C.

50. Add 20 µl resuspension solvent to the tube containing dried lipids, pipetting up and down along the sides.

**▴CRITICAL STEP** Some LCs have a maximum injection volume of 10 μl. Choose an appropriate volume that ensures complete resuspension without diluting the samples while accommodating maximum injection volume.

51. Vortex for 10 minutes in the cold room.

52. Sonicate for 5 minutes in an ice-cooled water bath.

53. Centrifuge for 10 minutes at 10,000 g in a pre-cooled centrifuge at 4°C. Transfer resuspended lipids to an appropriate LC-MS vial (Thermo Scientific, cat. no. 6PMCK30LVW). Avoid any particulate that might have centrifuged down at the bottom of the tube.

**▲CRITICAL STEP** Ensure that there are no bubbles at the bottom of your vial. Remove bubbles, if any, by gently tapping/flicking the vial with your index finger.

54. Load the vials into LC unit (cooled between 4°C and 8°C if possible).

55. Ensure the MS is properly tuned before running the sample. Types of tuning and their frequency will widely vary across MS instruments and detector types. Both mass accuracy and resolution are critical for proper lipid identification, as many lipids exist in similar retention times and *m/z* regions. Most MS tuning checks also involve a sensitivity check to ensure detection sensitivity. Perform these tunes following instrument specific protocols supplied by the vendor.

56. Inject 3 μl of this mixture from the cooled autosampler onto an Agilent Poroshell 120 (EC-C18, 2.1 x 100 mm, 2.7 μm, 1000 bar) column heated at 60 °C within the column compartment using an Agilent 1290 Infiniti II LC.

**▴CRITICAL STEP** Column heating allows for better retention time profiles through decreasing viscosity and increased lipid diffusion across mobile and stationary phases.

57. Use Mobile phase A [acetonitrile:H_2_O (60:40, vol/vol) supplemented with 0.1% formic acid and 7.5 mM ammonium acetate] and Mobile phase B [Isopropanol:acetonitrile (90:10, vol/vol) supplemented with 0.1% formic acid and 7.5 mM Ammonium Acetate] to separate peaks for detection on an Agilent Quadrupole Time-Of-Flight 6546 mass spectrometer.

**▲CRITICAL STEP** Use a Hamilton syringe for the formic acid. Use of plastic on highly concentrated acids can leach plasticizers which can contaminate LC-MS detection.

58. Program the LC method using the following gradient: starts at 85% A (15% B), decrease to 70% A over 2 minutes, then to 52% A over 30 seconds. Continue the gradient to 18% A over 12.5 minutes, followed by 1% A in 1 minute, and maintain this composition for 4 minutes. Return the gradient to 85% A and equilibrate the column for 5 minutes before next injection.

**▲CRITICAL STEP** We use this gradient that is optimized for global lipidome in our instrumental set up. For targeted or class-specific approaches or different instrument setups, this gradient can be modified.

59. Acquire mass spectra at the following parameters: *m/z* range: 300-1200, Gas Temp 250 °C, Gas Flow 12 L/min, Nebulizer 40 psi, Sheath Gas Temp 350 °C, Sheath Gas Flow 11, VCap (positive and negative mode: 4000 V), Nozzle Voltage 500 V, Fragmentor 175 V, Skimmer 65 V, Octopole RF Peak 750 V. Set MS/MS to trigger based on top 3 spectra/scan sorted by abundance with 2 spectra acquired per *m/z* prior to an active 0.2 min exclusion window. This data-dependent selection of precursor ions based on abundance provides a DDA-based MS approach. Set MS/MS acquisition threshold to an absolute value of 2500 or 0.01% of relative abundance. Set MS/MS scan range to 50–1000 *m/z* with a fixed collision energy of 25 eV. Perform acquisition in both positive and negative modes.

**▴CRITICAL STEP** The acquisition parameters described above were optimized for our instrumental setup, which is an Agilent 6546 Q-TOF coupled to an Agilent 1290 Infinity II LC. Depending on the LC-MS platform, detector type, ion optics, fragmentation mechanism, and sample complexity, optimization of parameters such as mass resolution, collision energy, and precursor ion selection may be required to achieve reliable lipid identification and quantification.

Data Analysis ●TIMING 2-6 h per batch

60. Open the LC-MS data in MS-DIAL and select the appropriate processing parameters according to the instrument configuration. Process the data using an appropriate lipid database (a subset of the LipidBlast *in silico* database was used in this study). More information regarding the use of MS-DIAL and comprehensive tutorial can be found at https://systemsomicslab.github.io/^41^. Review the processed MS-DIAL results to ensure that:

(i) Peak integration is consistent across all samples.

(ii) Fragmentation patterns are consistent with class- and species-specific library information.

(iii) Retention times are consistent with the expected chromatographic elution order for each lipid class.

61. Analysis of the standards can be performed using deuterated libraries. While MS-DIAL already has some embedded in the library, users can also build their own.

62. Use the exported output from the lipidomics ID software, MS-DIAL in this case, as an input to the Python script, available at https://github.com/guptaMSlab/Lipidomics-Python-Quant-Protocol^43^. For other software, ensure the exported output is compatible with the Python script – and ensure Python and Pandas are installed in the local computer.

**▲CRITICAL STEP** The scripts are compatible with the MS-DIAL default export format. No columns need to be added or deleted. For alternative software, ensure output follows template guidelines and begins on row 4.

63. Prepare two additional excel sheets (templates provided).

● One (template: lipidclass_RT) with three columns: labeled ‘Lipid Class’, ‘RT_low’ and ‘RT_high’. Populate the sheet with the lipid classes to be analyzed and corresponding minimum (RT_low) and maximum (RT_high) retention times for each lipid class.

**▲CRITICAL STEP** This allows for filtering of lipid classes based on established class-specific retention time windows to help reduce false positives. The units should match those in lipid software, and lipid classes should appear according to LIPID MAPS classifications used across most software. RT_low establishes the minimum retention time for that lipid class and RT_high establishes the maximum retention time for that lipid class.

● A quantification sheet (template: Template_Quantification_Information) with Standard names, Ontology, and Adduct types and concentration in each sample. Standard names should match those on your standard area sheet. Ontologies can be added or removed for that standard via the list separated by comma. The Standard concentration in each sample can be calculated as the moles injected onto the LC, assuming 100% recovery.

**▲CRITICAL STEP** Ensure sample names are exactly as they appear in the standard and lipid area sheets and followed by ‘standard conc’.

64. Run the Python script. When prompted, enter all sample names by copying them directly from the lipid area sheet. Columns containing ‘Blank’ in the name are excluded from quantitation, but selected blanks can be chosen for the next prompt to calculate noise thresholds for troubleshooting your LC-MS. The script generates output files containing unquantified sums of the area, quantified data, and an additional Excel file of quantified lipids separated by ontology. In addition, for troubleshooting, users can see how many lipid species fall below the threshold they set or exceed a set-fold over the average of the blanks.

### Troubleshooting

Troubleshooting advice can be found in Table 3.

### Timings

**Induction of LDs in cultured neurons: Day 1**

Steps 1-8, Primary culture of Cortical neurons: 6 h hands-on time

Step 9, Induction of LDs in cortical neurons: 1 d (30 min hands-on time)

*Lysis of neurons and purification of LDs: Day 16*

Step 10-14, Disruption of cortical neurons: 1.5 h hands-on time

Step 15-16, Preparation of PNS: 15 min hands-on time

Step 17-22, Purification of LDs by density-based floatation: 3 h (1 h hands-on time)

*Quality control: Day 17*

Step 23-24, Dynamic light scattering: 1 h hands-on time

Step 25-28, Confocal micrography: 1 h hands-on time

Step 29-36, Thin layer chromatography: 2 d (6 h hands-on time)

*LC-MS Data acquisition and analysis: Day 20*

Step 37-48, Extraction of lipids for LC-MS: 1 d (1.5 h hands-on time)

Step 49-59, DDA based LC-MS data acquisition: 6 h hands-on time

Step 60-64, Data analysis: 2-6 h hands-on time per sample

## ANTICIPATED RESULTS

Our step-by-step protocol (Fig. 1) facilitates reproducible induction, isolation, and characterization of NLDs from primary neurons. Following inhibition of neuron-specific triglyceride lipase (DDHD2) by KLH45, neurons display a clear increase in BODIPY-positive puncta in the soma as well as neurites relative to untreated controls (Fig. 2b). Electron microscopy reveals that the LDs are positioned near clusters of synaptic vesicles at the synaptic boutons (Fig. 2c), supporting the physiological relevance of these organelles at synapses^24^. Moreover, accumulation of TGs upon dose-dependent inhibition of DDHD2 lipase in neurons is also confirmed by TLC analysis (Fig. 2d,e).

The NLDs are part of a complex intracellular organellar milieu, and their separation, therefore requires sequential purification steps followed by quality check. Following cell lysis by nitrogen cavitation and flotation through a four-layer sucrose gradient (Fig. 3a,b), the top layer of the gradient (Fig. 3c) should be enriched with buoyant LDs that stain positive for both BODIPY and CellMASK staining neutral-lipid core and phospholipid covering respectively (Fig. 4a, left) In contrast, the lower layers of sucrose gradient are enriched with CellMASK positive bilayered membranous organelles but little or no BODIPY-positive puncta (Fig. 4a, right), indicating minimal loss of NLDs during floatation. Dynamic light scattering is expected to show a heterogeneous LD population, typically ranging from ∼200 nm to several micrometers (Fig. 4b). TLC analysis of the isolated NLD fraction shows strong enrichment of glycerolipids and sterol esters, while polar lipids remain at the starting point of the TLC (Fig. 4c,d). Excess phospholipid signal or diffused smearing would indicate membrane contamination and should prompt optimization of lysis or layering of sucrose gradient in steps 17-19 (Fig. 3b,c).

In LC-MS, users should expect robust detection of neutral lipids together with lower-abundance phospholipids, ether lipids, and other signaling lipids, provided that lipid extraction and resuspension are efficient (Fig. 5a). The class-level profile (Fig. 6a) should show enrichment of triglycerides in isolated NLD fraction relative to the whole neuron sample consistent with the neutral-lipid core of lipid droplets. At the same time, the detection of phosphatidylcholine and other phospholipids in NLD fraction is expected, as these monolayered lipids surround the hydrophobic core of the LDs. The species-level distributions provide a more detailed view of the major lipid classes and can be used to compare the acyl-chain composition of triglycerides and phosphatidylcholine between NLDs and whole-neuron input (Fig. 6b,c).

Together, the lipidomics data should be interpreted as evidence that the isolated fraction is enriched in a compositionally distinct LD population rather than representing a random subset of total neuronal lipids. Conversely, weak triglyceride enrichment, poor recovery of lower-abundance lipid classes, or highly similar profiles between NLDs and total neuron input may indicate incomplete floatation, sample loss, or inefficient lipid extraction in pervious steps.

Overall, successful implementation of this protocol will enable isolation and characterization of NLDs across diverse neuronal subtypes and models of neurodegenerative diseases. More broadly, this protocol provides a useful framework for advancing lipid research in neurobiology, and offers an entry point for investigating the complexity of NLDs in neural contexts and their potential relevance to neurological disorders.

## Data availability

The data, code, protocols, and key lab materials used and generated in this study are listed in a Key Resource Table alongside their persistent identifiers at https://doi.org/10.5281/zenodo.1966819877 and https://doi.org/10.5281/zenodo.2057514078. The raw lipidomics files have been deposited to MassIVE repository (Data set identifier: MSV000099675)^79^. All data cleaning, preprocessing, analysis, and visualization was performed using ImageJ/FiJi software and Python. An earlier version of this manuscript was posted to BioRxiv on Dec. 14, 2023, at DOI: https://doi.org/10.1101/2023.12.13.57152780.

## Code availability

All Python scripts used for the lipidomic analysis and database generation can be found at GitHub repository: https://github.com/guptaMSlab/Lipidomics-Python-Quant-Protocol and were given a persistent identifier via Zenodo: https://doi.org/10.5281/zenodo.2057514078.

## Acknowledgements

We thank Yumei Wu and Pietro De Camilli for providing EM images of synaptic terminal; and Anoop N. Pillai, Grace Dearden and Anant Menon for providing lab facilities for TLC analysis.

## Author Contributions

MK, KG, and TAR conceptualized the project. MK, RM, SR, and JAK performed the experiments and analyzed data. MK prepared the manuscript with help of other authors.

